# Genomic evolution reshapes cell type diversification in the amniote brain

**DOI:** 10.1101/2024.06.24.600323

**Authors:** Duoyuan Chen, Zhenkun Zhuang, Maolin Huang, Yunqi Huang, Yuting Yan, Yanru Zhang, Youning Lin, Xiaoying Jin, Yuanmei Wang, Jinfeng Huang, Wenbo Xu, Jingfang Pan, Hong Wang, Fubaoqian Huang, Kuo Liao, Mengnan Cheng, Zhiyong Zhu, Yinqi Bai, Zhiwei Niu, Ze Zhang, Ya Xiang, Xiaofeng Wei, Tao Yang, Tao Zeng, Yuliang Dong, Ying Lei, Yangang Sun, Jian Wang, Huanming Yang, Yidi Sun, Gang Cao, Muming Poo, Longqi Liu, Robert K. Naumann, Chun Xu, Zhenlong Wang, Xun Xu, Shiping Liu

## Abstract

Over 320 million years of evolution, amniotes have developed complex brains and cognition through largely unexplored genetic and gene expression mechanisms. We created a comprehensive single-cell atlas of over 1.3 million cells from the telencephalon and cerebellum of turtles, zebra finches, pigeons, mice, and macaques, employing single-cell resolution spatial transcriptomics to validate gene expression patterns across species. Our study revealed significant species-specific variations in cell types, highlighting their conservation and diversification in evolution. We found pronounced differences in telencephalon excitatory neurons (EX) and cerebellar cell types between birds and mammals. Birds predominantly express *SLC17A6* in EX, whereas mammals expressed *SLC17A7* in neocortex and *SLC17A6* elsewhere, possibly due to loss of *SLC17A7* function loss in birds. Additionally, we identified a novel bird-specific Purkinje cell subtype (SVIL+), implicating the LSD11/KDM1A pathway in learning and circadian rhythms, and related numerous positively selected genes in birds, suggesting an evolutionary optimization of cerebellar functions for ecological and behavioral adaptation. Our findings elucidate the complex interplay between genetic evolution and environmental adaptation, underscoring the role of genetic diversification in the development of specialized cell types across amniotes.

## Introduction

Amniotes, which evolved over 320 million years ago (MYA), including reptiles, birds, and mammals, are among the most abundant and widely distributed terrestrial animals. This transition from aquatic to terrestrial habitats accelerated the evolution of amniote brain information-processing capabilities, thereby enhancing adaptation to terrestrial environments (Woych et al., 2022). Extensive research has revealed significant variations in the morphology and connectivity of the brain throughout the evolution of amniotes (Belgard et al., 2013; La Manno et al., 2016; Zeisel et al., 2015). Among the amniote taxa, avians represent an independently differentiated branch, characterized by highly specialized brains. Relative to body weight, the avian brain is notably large, and features complex structures and functions (Ksepka et al., 2020; Sayol et al., 2019). The avian brain uniquely exhibits higher neuron and synapse densities compared to other animal groups, as well as distinct telencephalic structure and connectivity patterns relative to mammals (Herculano-Houzel, 2020; Sayol *et al*., 2019). The distinctive structure and functionalities of the avian brain are intrinsically linked to their unique survival strategies, serving as a crucial neural foundation for adaptation and reproduction in dynamic and complex environments. In contrast, understanding the mechanism underlying these distinctive structures and functionalities reveals a significant gap between complex function and cellular or molecular changes. Gene expression datasets have been employed to address the challenges of comparative analysis for the evolutionary trajectories of brain cell types across multiple species (Belgard *et al*., 2013; He et al., 2017; Tosches et al., 2018).

Recently, single-cell RNA sequencing has emerged the prominent technique for detailed comparative investigations of cell type composition (Gumnit and Tosches, 2023; Hain et al., 2022; Shafer et al., 2022; Styfhals et al., 2022; Tosches *et al*., 2018; Woych *et al*., 2022). Through comparative analysis, we found that gene family evolution and shifts in paralog expression contribute to cellular diversity (Shafer *et al*., 2022). Comparative single-cell sequencing research in birds and mammals has predominantly focused on well-defined brain subregions (Bakken et al., 2021; Colquitt et al., 2021; Franjic et al., 2022; Ma et al., 2022). Furthermore, studies have generally targeted a narrow range of closely related species (Wang et al., 2021). The absence of a comprehensive single-cell atlas of the avian brain limits our detailed understanding of its cellular specificities. Moreover, investigations into more diverse and distantly related species, crucial for understanding the evolutionary dynamics of cell types and brain functions, remain scarce. Challenges in this field partly stem from the absence of a unified data platform and standardized technologies to produce cross-species datasets. Additionally, the field lacks robust integration analysis methods to comparing single-cell data across distantly related species.

Using a standardized single-nucleus RNA sequencing (snRNA-seq) platform DNBelab C4 (Liu et al., 2019), we constructed a comprehensive cell-type atlas for reptiles (turtles), two birds (pigeons and zebra finch), and mammals (macaques and mice). We supplemented this with spatial transcriptome analyses using Stereo-seq (Chen et al., 2022a) for several critical brain regions to validate our findings. Our global analysis revealed that significant functional divergence or loss of paralogous genes has driven the evolution of brain cell types. Notably, differences in cortical excitatory neurons between birds and mammals were primarily influenced by the expression of the paralogous genes *SLC17A6* and *SLC17A7*, which correlate with genomic variations between species. Detailed structural analyses of the transmembrane proteins encoded by these genes revealed that minor mutations could cause substantial changes in their transmembrane domains. We have identified a newly emerged, rapidly evolving Purkinje cell type (SVIL+) in birds, alongside significant changes in Granule and Bergmann cell types. These changes in expression patterns may indicate how the avian cerebellum has adapted to new behavioral lifestyles, such as flight. These findings suggest that these cell types are crucial for adaptation to unique ecological niches and the demands of aerial mobility in birds. Our results highlight a correlation between genomic evolution and cell type differentiation in amniotes, supporting the theory that genomic evolution through natural selection is crucial for the evolutionary differences in brain function. This resource is interactively accessible at: https://db.cngb.org/cdcp/dataset/SCDS0000639/.

## RESULTS

### A cell-type atlas of the amniote brain

To better understand the evolution of cell-types in the amniotie brain, we constructed a comprehensive single-cell atlas using a droplet based DNBelab C4 snRNA-seq (Liu *et al*., 2019) across species, complemented by several sections of spatial transcriptomic (Stereo-seq) data for validation (Chen *et al*., 2022a). This included three newly generated datasets in this study from the Sauropsida branch: Chinese soft-shell turtle (*Pelodiscus sinensis*), zebra finch (*Taeniopygia guttata*) (hereafter referred to as ‘finch’), and pigeon (*Columba livia*). Additionally, we incorporated published data from two mammals: mice (*Mus musculus*) and cynomolgus monkey (*Macaca fascicularis*) (Bakken *et al*., 2021; Kozareva et al., 2021; Yao et al., 2021) **(Figure 1A)**. Given the complex architecture of avian brains, we conducted snRNA-seq with detailed parcellation in zebra finches and pigeons, encompassing the pallium, subpallium, optic tectum (OT), cerebellum, thalamus, midbrain, and hindbrain (here, ‘hindbrain’ includes the pons and medulla but excludes cerebellum here). The avian pallium was subdivided into the hyperpallium, mesopallium, arcopallium, nidopallium, and nidopallium caudolaterale (NCL) **(Figure 1B**; **Table S1)**.

**Figure 1.**
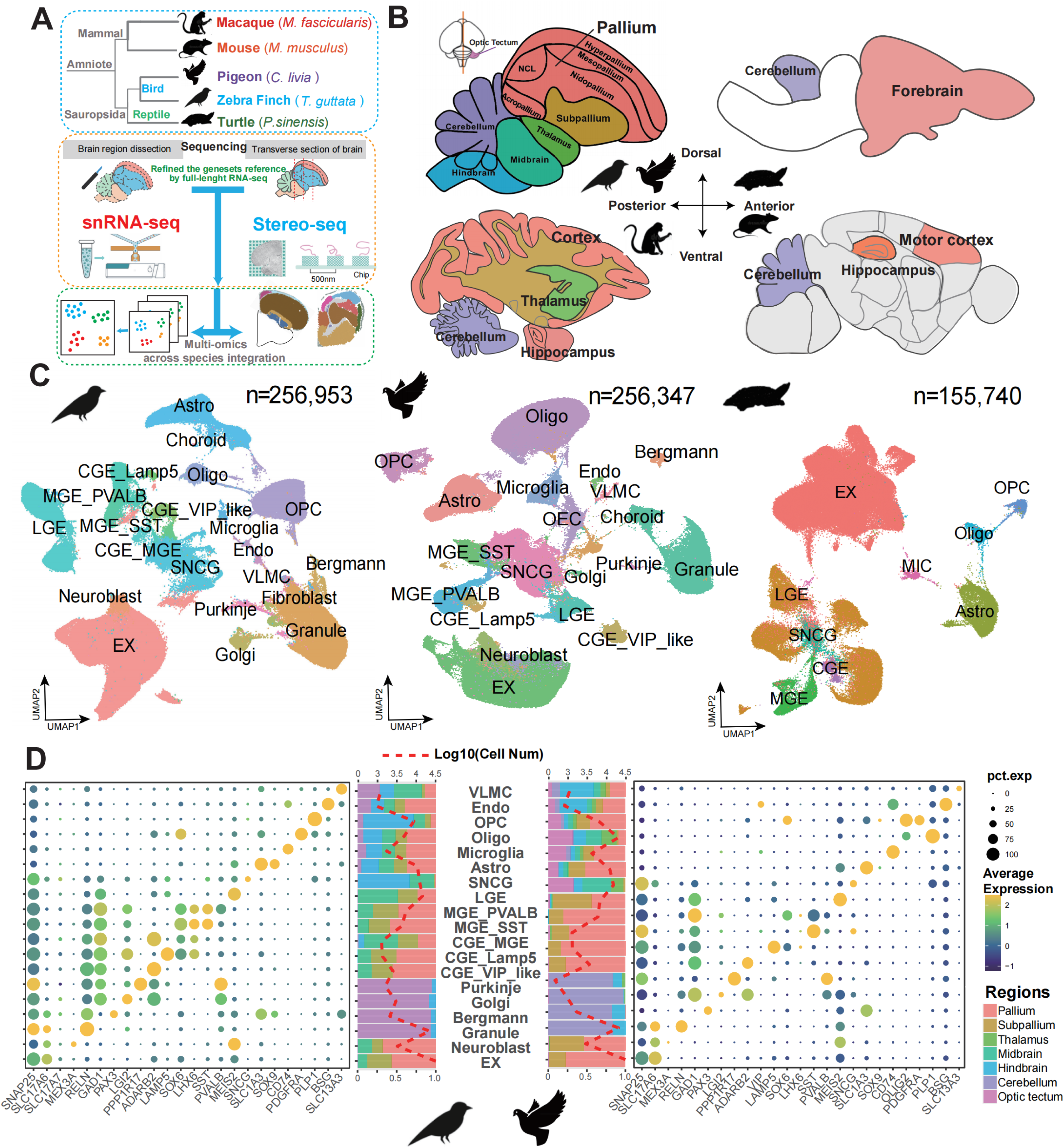
Cell-type conservation and diversity in amniotic brain. **(A)** Schematic diagram of the complete design analysis model in this paper. Species selection: Sauropsida branch: Chinese soft-shell turtle (Pelodiscus sinensis), zebra finch (Taeniopygia guttata), and pigeon (Columba livia). mammals: mice (Mus musculus) and cynomolgus monkey (Macaca fascicularis); Multi-omics sequencing; Integration and comparison across species. **(B)** Brain regions sampled in detail for snRNA-seq with all the five species. For the pigeon and finch, the brain was divided into 11 areas, which were integrated into 7 major areas in subsequent analyses, including pallium, subpallium, optic tectum (OT), cerebellum, thalamus, midbrain, and hindbrain (‘hindbrain’ includes the pons and medulla but excludes cerebellum here). The pallium was subdivided into hyperpallium, mesopallium, arcopallium, and nidopallium caudolaterale (NCL). **(C)** UMAP of single cell atlas of the brain derived from pigeons, finches, and turtles, coloured by cell type. Some cell type names were abbreviate: excitatory neurons (EX), astrocyte (Astro), oligodendrocyte precursor cells (OPC), oligodendrocytes (Oligo), endothelial cells (Endo) and vascular leptomeningeal cells (VLMC). Since there is no well-defined VIP gene in birds, the cell type homologous to the mammalian CGE_VIP was named CGE_VIP_like. **(D)** Dotplot depicting the expression of marker genes across cell types, with the proportion of cell sampling sources represented by variably colored bars. The dashed line indicated the logarithm of the total number of cells for each cell type.

To improve the gene annotation quality in turtles and pigeons, we aligned the raw reads to the reference genomes, integrating a refined gene set annotated via bulk RNA-seq of brain tissue using long-read sequencing such as PacBio (Rhoads and Au, 2015) and Cyclone (a new third-generation sequencing platform from BGI) (see **STAR Methods**). We increased the total number of annotated genes from 24,856 to 25,892 in turtles and from 15,392 to 19,201 in pigeons, and enhanced the completeness of gene structure, updated with a complete gene structure with 97% of genes in turtles and 98% in pigeons in the refined gene sets. Using these refined gene sets as a reference, we significantly improved mapping efficiency, with the proportion of reads aligned to gene regions increasing from approximately 45% to 75% for both species. This enhancement led to an increase in the number of genes detected per cell from approximately 800 to 1,000, and in unique molecular identifiers (UMIs) per cell from about 1,200 to 1,800. After quality filtering, we obtained a total of 1,391,446 cells, of which 669,040 cells were generated in this study for the following species: turtles (155,740), pigeons (256,347), and finches (256,953) (**Figure 1C**; **Table S1)**.

To annotate the cell types across species, we refined the single-cell graph convolutional neural network model (scGCN) (Song et al., 2021), which we termed ‘entropy-scGCN’ (see **STAR Methods)**. By employing the mouse as a reference, we considered several factors for annotating cell types in other species: (1) To including more homologous genes in the amniotes with significant speciation time in the subsequent analysis, we calculated the gene expression of the homologous genes within gene families **(Table S2)**, extending beyond simple one-to-one orthologous relationships; (2) To incorporate entropy theory to evaluate cell type diversity. We identified 19 main interspecies cell types by applying Louvain clustering followed by the entropy-scGCN approach (**Figure 1C**). The expression patterns of well-established marker genes aligned with expectations and demonstrated high conservation across species (**Figure 1D**). (3) Based on the individual clustering by each species and differential gene expression patterns, we subdivided the major cell types into further subtypes (24 in turtle, 45 in pigeon, and 48 in finch) (**Figures S1A and S1B**).

In conclusion, we have constructed a single-cell atlas of amniote brains using a consistent technology platform and generated datasets for unbiased cross-species comparisons.

### Integration and comparison across amniotes

The identification of homologous cell types across species is crucial for understanding cell type evolution and is influenced by various factors. Traditionally, conserved cell types across species have been defined by canonical marker genes, as demonstrated in model species such as primates and rodents (Chen *et al*., 2022a; Han et al., 2022; Qu et al., 2022; Wang *et al*., 2021). However, this approach frequently encounters significant limitations, particularly for non-conserved cell types and when addressing a broader range of species. Therefore, it has been proposed that using the historical continuity of gene regulatory networks rather than the expression of individual homologous genes will be more effective (Wagner, 2007). For comparative analysis across multiple species, especially those that diverged long ago, employing comprehensive gene information for identification is particularly crucial. To address this issue, as previously described, we primarily aligned homologous genes across species based on gene family information (see **STAR Methods**). To minimize biases introduced by varying cell counts, we conducted integrated clustering of single cells using downsampling to ensure equality across five species, totaling 195,162 cells. We conducted scGCN all-against-all species comparisons (**Figure S2B**) and detected recognized cell marker genes, identifying 25 major cell types (**Figure 2A**) and 63 cell subtypes (**Figures 2D and S2A**). As expected, conservation among major cell types was significantly higher than among subtypes, indicating that interspecies differences in cell types are primarily manifested in the differentiation of cell subtypes (**Figures 2A, 2D, and S2F**). Particularly in excitatory neurons, where almost all cell subtypes displayed clade-or species-specific characteristics (**Figures 2D and S2F**).

**Figure 2.**
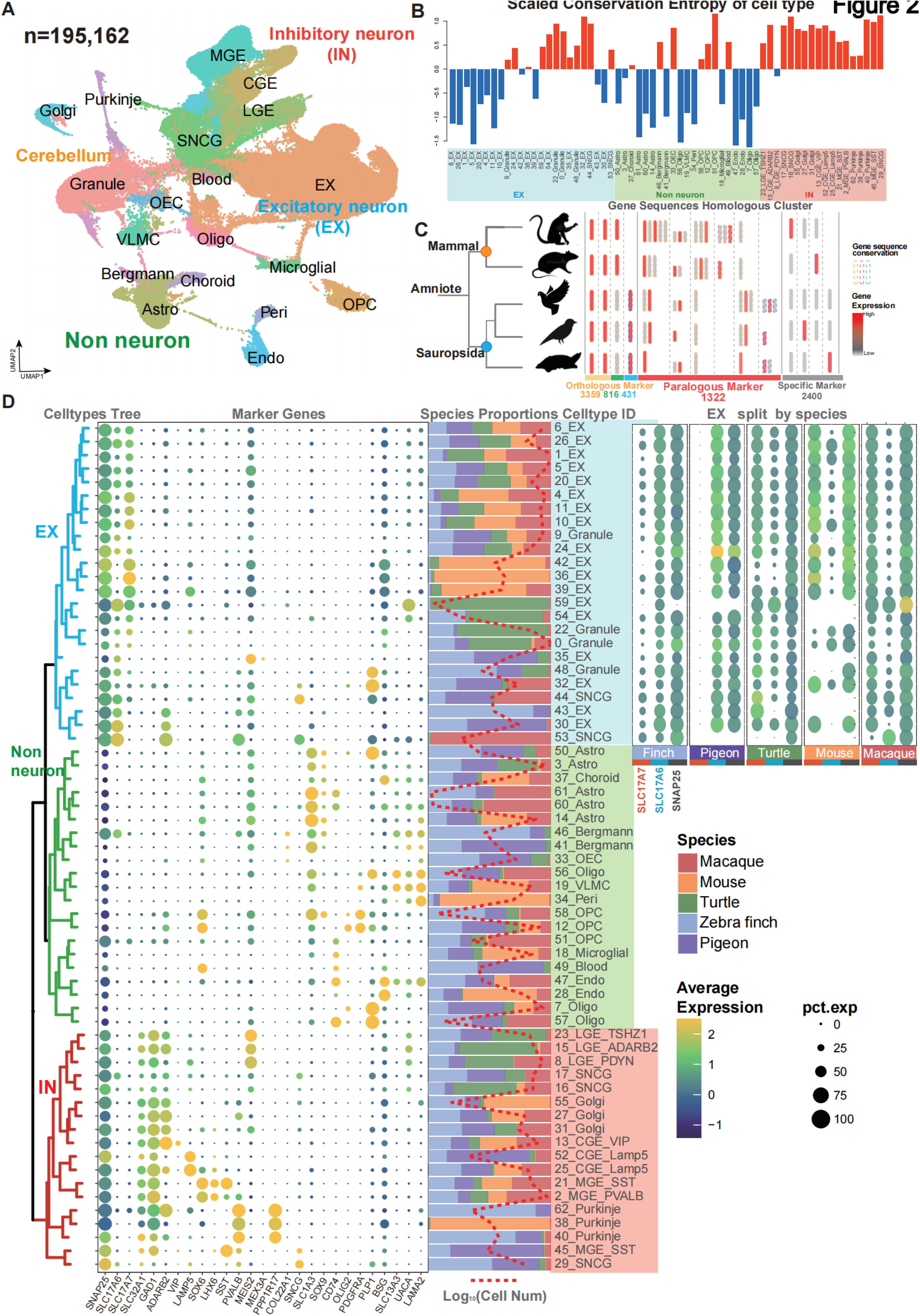
Integration and comparison of the single cell atlas across amniotes. **(A)** Integrated cross-species UMAP atlases of the whole brain, downsampling to approximately ∼40,000 cells per species **(Table S1)** covering five species. On the basis of FIG. 1D, 3 additional cell types were finely differentiated: pericytes (Peri), olfactory ensheathing cells (OEC) and Choroid cells. **(B)** Scaled conservation entropy values for 63 cell type clusters across species, determined using the scGCN algorithm. Sub cell-type with larger values indicate significant differences. **(C)** Based on the characteristics of cell type marker gene families, gene expression was classified into three categories: one-to-one conservation across species, homologous gene family conservation, and species-specific. The schematic plot displays the classification and statistics. The number of genes for each type is indicated below the corresponding color. **(D)** Overview of cell taxonomy, marker gene expression, and cell-type abundance across species. Left: Cell taxonomy of 63 cell clusters, framed by color according to cell type categories including excitatory neurons (EX, blue), inhibitory neurons (IN, red), and non-neuronal cells (non-neuron, green). Middle: Dot plots illustrate the expression of marker genes used for annotating cell clusters shown in (A), where dot size and color represent the proportion of cells expressing each gene and the mean expression level within each cluster, respectively. Right: Percentage bar plots showing the proportion of 5 species in each cell type. The red dashed line represents the logarithm of the cell number. Top right: EX cell types were split into five species sources. Dot plot shows the expression of *SLC17A7*, *SLC17A6* and *SNAP25* genes.

To deeply investigate the diversity of cell subtype s across species and identify the emergence of new sister cell subtype s, we calculated information entropy values based on the all-against-all species scGCN mapping of each cell subtype (**Figure S2B**, see **STAR Methods**). A low entropy score for a specific cell type indicated that it was conserved and could be uniquely mapped to a corresponding cell type in other species via scGCN, signifying evolutionary conservation across species. Conversely, a high entropy score suggested a complex many-to-many mapping relationship across species and predicted the emergence of new sister sub-cell types (**Figure S2B; Table S4**). We observed that EX groups with low entropy and conserved types, such as 5_EX and 11_EX, exhibited relatively uniform proportions across species. In contrast, EX groups with high entropy that varied across species either showed a biased species proportion (22_Granule and 54_EX) or divergent gene expression patterns (e.g., 24_EX expressed *SLC17A6* and *SLC17A7* in birds and mammals respectively) (**Figures 2B and 2D**). Most non-neuronal types displayed conserved low entropy, with exceptions including premature oligodendrocytes (OPC and OEC) and cerebellar Bergmann cells (**Figure 2B**). To our surprise, most inhibitory neuron groups (IN) followed a high entropy pattern, although these cell types exhibited no significant biased proportion across species (**Figures 2B and 2D**). Consequently, we proposed that IN cells exhibit diverse gene expressions patterns and might be subdivided into more species-specific subtypes, a hypothesis we confirmed in subsequent analyses (**Section** regarding the IN groups).

Furthermore, based on the distribution of cell-type proportions, we identified several clade-enriched cell subtypes, specifically in birds or mammals (**Figure 2D**). Most excitatory neurons, including three bird-enriched types (30_EX, 35_EX, 43_EX) and all mammal-enriched EX types (36_EX, 39_EX, 42_EX), along with certain inhibitory and non-neuronal cells in the cerebellum (38_Purkinje, 40_Purkinje, 62_Purkinje, 41_Bergmann, 46_Bergmann), exhibited significant clade enrichment. Notably, by analyzing species-specific gene expression, we identified two distinct groups of EX cell types: avian-specific and mammal-specific. Avian EX cells predominantly express *SLC17A6*, while mammalian EX cells primarily express *SLC17A7*, with a minority of subtypes also expressing *SLC17A6*, and turtle EX cells expressing both genes (**Figure 2D**).

In the integrated dataset, we identified sub-cell types with unique expression patterns in specific clades or species. Therefore, we decided to explore how genes contribute to this evolutionary progress. To investigate the gain, loss, expansion, or contraction of genes or gene families contributing to cell type divergence across species, we first categorized the gene families of five species into types that were conserved and distinct for each branch (**Figure 2C**), then we identified cell type marker genes for each species independently; finally, we compared the expansion or contraction of gene families with marker genes across clades (**Figures 2C and S2C, Table S3**). Using this method, we categorized the marker genes into three categories (see **STAR Method**): orthologous markers (55% of all marker genes, with 40% conserved across all amniote species), paralogous markers (16%), and species-specific markers (29%) (**Figures 2C**). We calculated the conservation of cell type marker gene families across species (see **STAR Methods**), finding that the proportion of gene families conserved across all amniote species or clades (33%) was significantly higher than that of orthologous markers (19%) **(Figures S2E)**.

Furthermore, we compared transcription factor (TF) marker genes (Shen et al., 2023) with positively selected genes as reported in the literature (Warren et al., 2010) (**Tables S5 and S6**). The results indicated that, compared to effector genes (the remaining genes when excluding TF and positively selected genes (PSG)), TF genes exhibit greater conservation across species, while positively selected genes display the most pronounced differential expression (**Figure S2D**). These results suggest that conserved orthologous genes contribute to the uniform gene expression pattern observed across species. It is noteworthy that paralogous genes also play a significant role in shaping expression patterns across species, suggesting that different species may use various paralogous genes to maintain conserved cell types, as exemplified by *SLC17A6* and *SLC17A7*, specifically expressed in birds and mammals, respectively (**Figure 2D**). Taken together, the integrated cross-species single cell atlas enables us to accurately identify both conserved and diverse cell types, and to explore how variations within gene families are associated with and influence changes in these cell types.

### Excitatory neurons in the cortex of amniotes

Excitatory neurons exhibit increased cellular heterogeneity brain region specificity, and are more prevalent within specific clades in terms of sub-cell types, as previously described (**Figure 2D**). For systematic characterization and comparison, we isolated EX neurons from the pallium regions, specifically the pallium in birds, cortex in macaques, primary motor cortex in mice, and forebrain in turtles, for comparative analysis **(Figures 3A-D and S3A)**. We observed that similar cell types cluster together rather than by species, indicating an elimination of batch effects. However, when examining specific subtypes, we observed in the UMAP results that cells from phylogenetically closer species tend to cluster together (**Figure 3B**). Notably, several avian cells cluster together with the upper layers (L2-3) and deeper layers (L5-6) of the cortex of both macaques and mice (**Figures 3C and 3D**), suggesting that birds, like mammals, also harbor EX cells specialized for short- and long-range connections, respectively. Moreover, a significant spatial pattern was observed for avian EX cells, distinct from the laminar structure typically found in mammalian neocortex (**Figures S3B and S3C**). Furthermore, L4 cells in mammals did not have a significant counterpart in birds, indicating that L4 EX cells may be specific to mammals **(Figures 3C and 3D)**.

**Figure 3.**
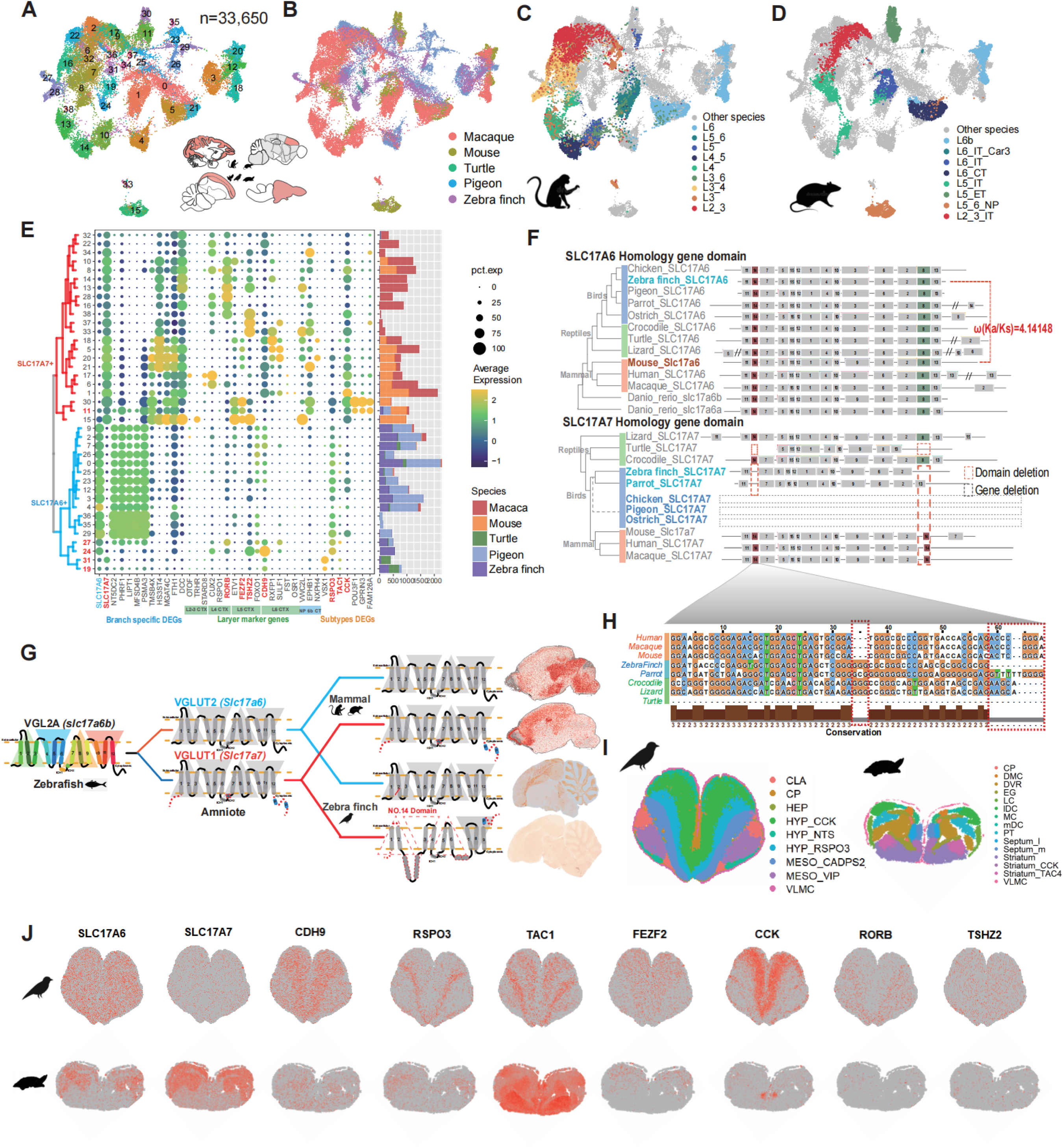
Evolutionary differences in excitatory neurons in the cortex and subcortex of amniotes. **(A-D)** UMAP plots of re-clustered excitatory neurons from the cortical areas of five species, with an integrated UMAP for all species shown in (A), color by species in (B), and EX neurons across cortical layers in macaques (C) and mice (D). **(E)** Expression of selected marker genes and the clustering relationships among all subtypes of EX neurons. Automatically clustered into two groups: SLC17A6+ (lower group in blue font) and SLC17A7+ (upper group in red font). Marker genes were selected based on clade- or species-specific expressions (blue), layer-specific marker genes in the neocortex (green), and DEGs of the two major groups (orange). **(F)** Gene tree illustrating the motifs of the *SLC17A6* (upper panel) and *SLC17A7* (lower panel) gene families in amniotes, detected using the MEME (Bailey et al., 2015). The location where the domains mutation occurred is marked with a red dashed line. Species without corresponding genes of the same name are marked by gray dashed lines. Mouse and finch *SLC17A6* genes were detected by Tree2gd (Chen *et al*., 2022b) with Ka/Ks significantly greater than 1, marked on the side. **(G)** Schematic diagram showing the structural changes in VGLUT2 (encoded by *SLC17A6*) and VGLUT1 (encoded by *SLC17A7*) proteins throughout the evolution of zebrafish, predicted in the common ancestor of amniotes, mammals, and birds. The right panel depicts the spatial distribution of gene expression (sagittal) in a brain section. **(H)** Difference sequence alignment diagram for the no.14 domain in (F) and the corresponding protein variations between species. Amino acid sites showing significant differences between birds and mammals are highlighted in red dashed boxes. **(I)** Brain domains of two sections from finches and turtles analyzed using the spatial transcriptomic (Stereo-seq)(Chen *et al*., 2022a) atlas (see Data availability). **(J)** Spatial patterns of gene expression for several marker genes (E) in two sections from (I).

SLC17A6+ and SLC17A7+ have been identified predominantly in avian-enriched and mammal-enriched EX neurons, respectively (**Figure 2D**). To delineate these two distinct clade-enriched groups of EX cell types, we clustered all 39 EX cell subtypes using canonical marker genes (e.g., *SLC17A6* and *SLC17A7*), layer-specific marker genes (e.g., *CUX2*, *RORB, CDH9*), and other species-specific genes (**Figure 3E**). As expected, we identified two distinct groups of cell types: mammal-specific and sauropsid-specific, characterized by *SLC17A7* and *SLC17A6* expression, respectively. Notably, we identified several genes with expression patterns similar to those of the two canonical genes. Five genes, namely *TMSB4X, HS3ST4, MGAT4C, FTH1*, and *DCC*, were highly expressed in SLC17A7+ EX neurons, associated with functions of axon growth (*DCC*) (Duman-Scheel, 2009), neural circuit formation, and synaptic plasticity (*TMSB4X*). Additionally, cortex layer marker genes and five other genes (namely, *NT5DC2, PHRF1, LIPT1, MFSD4B,* and *PSMA3*), associated with functions of neuron energy metabolism (such as *LIPT1*) and neurotransmitter transport (such as *MFSD4B*) (Perland et al., 2017; Soreze et al., 2013), were found to be highly expressed in SLC17A6+ cells. Cluster 24 (**Figures 3E**), part of the SLC17A6+ cell group and primarily located in the finch brain, showed high expression of deep-layer markers (L5-6) such as *TSHZ2* (related to neural circuitry development) (Yao *et al*., 2021), *CDH9* (associated with neuronal connectivity and plasticity) (de Wit and Ghosh, 2016), and *RSPO3* (linked to synaptic plasticity and notably concentrated in the finch hyperpallium) (Lovell et al., 2008) (**Figures 3I and 3J**). Our findings suggested an evolutionary adaptation towards long-range synaptic connections and enhanced neural communication and plasticity (de Wit and Ghosh, 2016; Polanco et al., 2021). Interestingly, cluster 31, also within the SLC17A6+ group, unexpectedly expressed *SLC17A7* in turtles but not in finches, showed the expression of the deep-layer marker *CDH9*, similar to cluster 24 (**Figures 3E**). This marker is notably enriched in the hyperpallium of birds (**Figures 3I and 3J**), suggesting that sauropsids may retain some EX cells expressing *SLC17A7*, a trait typically specific to mammalian cortex. These findings suggested that EX neurons in sauropsids exhibit significant divergence and retain functions analogous to those of deep-layer, long-range connection neurons in mammals (Hussan et al., 2022).

To elucidate the evolutionary mechanisms underlying these two cell-type groups, we performed a comparative analysis on protein sequences and the protein domain folding. *SLC17A6* and *SLC17A7* encode the proteins VGLUT2 and VGLUT1, respectively, both involved in the transport of the neurotransmitter glutamate into synaptic vesicles (Park et al., 2019; Reimer, 2013; Serrano-Saiz et al., 2020). Utilizing the UniProt database (UniProt, 2023) to confirm protein structures, and integrating the gene family dataset, we reconstructed the evolutionary trajectory of these proteins from zebrafish to birds and mammals (**Figure 3G**). *SLC17A6* underwent a duplication event in the ancestral lineage of amniotes, resulting in the paralog *SLC17A7*. Motif sequence analysis of these two genes across amniotic species (**Figure 3F**) revealed that *SCL17A6* motifs, conserved across amniotes, experienced positive selection pressure in the bird clade (Ka/Ks=4.14 compared to mice). *SLC17A7* either lost the entire gene or specific motifs in birds and turtles, consistent with the observed lack of gene expression in pigeons and limited partial expression in finches.

Upon comparing the motifs of these two genes, we observed that in mammals, *SLC17A7* harbors a duplication of motif-14 positioned closer to the 3’ end compared to *SLC17A6*, whereas in finches, this motif was absent (**Figure 3F**). Multiple alignment of motif-14 revealed a conserved gene structure in mammals and divergence in sauropsids (**Figure 3H**). The protein structures (**Figures 3G and S3D**) elucidated the associations between structural mutation sites and the protein’s transmembrane domain. Variations in their intramembrane sequences have been demonstrated to influence their glutamate affinity and regulatory capabilities. Previous research has shown that *SLC17A6* is associated with higher glutamate concentrations, while *SLC17A7* exhibits faster response kinetics (Fremeau et al., 2001; Fremeau et al., 2004; Herzog et al., 2006).

Collectively, these findings suggested that through natural selection, specific characteristics are retained in mammalian species: *SLC17A6* is expressed in certain cortical cell types, more broadly in subcortical brain regions and the cerebellum, as well as areas with higher glutamate concentrations, whereas *SLC17A7* is expressed in the cortex to facilitate rapid response. Birds, however, exhibited overall higher brain glutamate concentrations than mammals, necessitating a long-term stable glutamate regulation mechanism, making *SLC17A6* more suitable for this role (Cheret et al., 2021; Sreedharan et al., 2010). Due to natural selection, some species, like chickens, gradually lost *SLC17A7*, while others like finches, retained non-synonymous mutations in *SLC17A7* **(Figures 3F and 3H)**, disrupting its structural domain and resulting in whole-brain expression of *SLC17A6* (**Figure 3G**). Overall, significant differences in gene expression exist between avian and mammalian cortical excitatory neurons. These disparities could be attributed to gene duplication, site mutations, and other genomic evolutionary events, suggesting that comparative genomics analysis could effectively elucidate single-cell data to highlight differences between cell types.

### Evolution of distinct cerebellar cell types and expression characteristics in Birds

The cerebellum, thought to have originated early in vertebrate evolution, is present across a broad spectrum of vertebrates, including fish, reptiles, and mammals (Butts et al., 2014; Hashimoto and Hibi, 2012; Hibi et al., 2017; Sugahara et al., 2017). However, the complexity and structure of the cerebellum have evolved variably across species (Kebschull et al., 2020; Sepp et al., 2024). To examine the specific cell types present in sauropsids and mammals, we isolated 200,311 cells from their cerebellum, identifying a total of thirteen distinct cell types (**Figures S4A and S4C)**. As expected, the granule cells constituted the majority of the cell population (**Figures 4A and S4B**). To further analyze cerebellum-specific cell types, we isolated an additional 174,056 cells, comprising granule, Bergmann, Golgi, and Purkinje cells, distinguished by their unique characteristics in the cerebellum compared to other brain regions (**Figure 4B**). Compared to other brain regions, cerebellar cell types, particularly Bergmann, Golgi, and Purkinje cells, showed greater inter-species variability (**Figures 2D and 4B**), indicating significant functional evolution within the cerebellum of amniotes to adapt to their environment. Specifically, *SOX2* and *SOX9* were predominantly expressed in mammalian and avian Bergmann cells, respectively (**Figure S4D**). Marker genes of granule cells, such as *GABRA1*, *GABRA4*, and *GABRA6*, demonstrated differential expression between birds and mammals (**Figure S4E**). Furthermore, genes including *PCLO, LGI2, NXPH2*, were notably expressed in the cerebellar cells of birds (**Figures S4D and S4F**), which were involved in formation and maintenance of synapses and the transmission of neural signals (Cen et al., 2008; Favuzzi et al., 2019; Petrenko et al., 1996).

**Figure 4.**
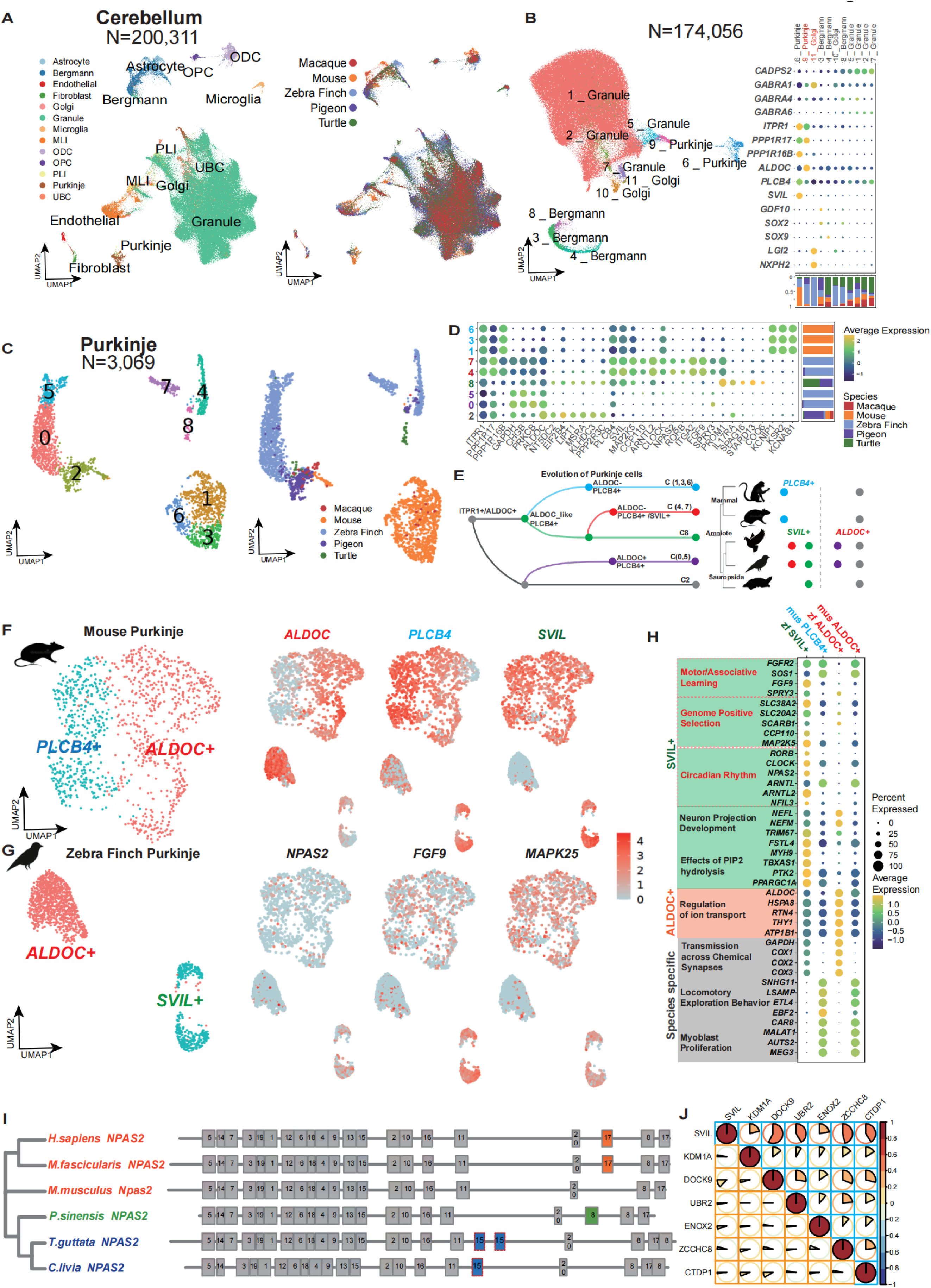
Cell type diversity in amniote cerebellum. **(A)** Clustered UMAP plot showing the cell annotation results of the entire cerebellum cells from 5 species. The species origin distribution is shown in the right. **(B)** UMAP of the clustering of cerebellar-specific cell types (granule cells, Bergmann cells, Purkinje cells, and Golgi cells) in 5 species. Dotplots of marker gene expression for each cluster are shown on the right. The proportion of species is indicated in the bar plot with the corresponding color. **(C)** Clustered UMAP plot showing Purkinje cell classification for 5 species. Subtypes numbers are shown on the left and species origin on the right. **(D)** Dot plot of conserved and differential gene expression in each Purkinje subcell-types (C). Species origin proportions are shown on the right. **(E)** Diagram of the inferred evolutionary pattern of Purkinje cells in amnioids, based on the conservation of cell subtypes. Right: distribution of the corresponding isoforms in each species, with color corresponding to (D). **(F-G)** Expression profiles of marker genes that show differential expression in the two subtypes of Purkinje cells between the mouse and finch. **(H)** DotPlot illustrating differentially expressed genes between the two subtypes of Purkinje cells in both species. Genes are categorized based on their function. **(I)** The circadian gene NPAS2, which is highly expressed in the SVIL+ group of zebra finch, domains divergence tree obtained by MEME (Bailey *et al*., 2015) between species. **(J)** Plot of the expression association between the SVIL gene and the downstream regulatory genes of LSD1 (KDM1A) in mouse (orange) and zebra finch (blue).

Notably, Purkinje cells, which exhibited the most pronounced species differences (**Figure 4B**), comprised two distinct subtypes: 6_Purkinje and 9_Purkinje, each characterized by the expression of specific markers, including *ITPR1, PPP1R17, ALDOC, PLCB4*, and *SVIL* (**Figure 4B**). We re-clustered the 3,069 Purkinje cells and identified nine cell subtypes, which were subsequently categorized into five groups based on species composition and UMAP distributions: C2 (all species), C8 (sauropsids), C(0,5) (birds-specific), C(4,7) (finch-specific), and C(1,3,6) (mammal-specific) (**Figures 4C and 4D**). We reconstructed the evolutionary history of Purkinje cell types across amniotes based on marker expression patterns (**Figure 4E**), identifying C2 as the common cell type among amniotes, and C(1,3,6) as the mammal-specific subtype (marked by *KCNIP1, KCNAB1, and KSR2* expression). Notably, C(0,5) (marked by *ALDOC* expression) and C(4,7) (marked by *SVIL* expression) were bird-specific but exhibited distinct expression patterns. As reported by Xiaoying Chen et al., two major Purkinje subtypes were identified in mice: one predominantly expressed *ALDOC* and the other *PLCB4,* findings that our study corroborates (Chen et al., 2022c). In contrast, birds also possessed ALDOC+ Purkinje cells, and also harbored a bird-specific subtype, SVIL+ Purkinje. Subsequent DEG analysis of these Purkinje subtypes revealed that ALDOC+ cells from both finch and mouse share conserved marker genes (*ALDOC, HSPA8, RTN4, THY1,* and *ATP1B1*) (**Figures 4F, 4G and 4H**) with association of regulation of ion transport, suggesting that ALDOC+ represents an ancestral Purkinje subtype in early cerebellar evolution.

To deeply investigate the SVIL+ Purkinje cells in birds, we focused on the up-regulated genes and their changing regulatory networks. SVIL+ cells were characterized by the expression of genes associated with circadian rhythms, including *RORB, CLOCK, NPAS2,* and *ARNTL* (**Figure 4H and S4G; Table S7**). Comparison with positively selected genes in birds revealed overlaps, including *SLC38A2, SLC20A2, CCP110*, and *MAP2K5*, suggesting significant positive selection pressure on the emergence of new SVIL+ Purkinje cells in birds (**Figure 4H**). We identified mutations in protein sequences, including motif domain gains or losses, were identified in bird SVIL+ cells. For example, motif alignment for the marker gene *NPAS2* across amniotes revealed a motif-15 gain in birds, an motif-8 duplication in turtles, and a motif-17 duplication in mammals (**Figure 4I**). In the functional enrichment of DEGs, the SVIL+ cells exhibited enriched functions related to circadian rhythms, neuron projection development, and effects of PIP2 hydrolysis (**Figures 4H and S4I**). Those results may indicate a significantly greater number of dendritic branches in birds compared to mammals, which was consistent with previous observations (Cunha et al., 2021; Kidd, 2017). We analyzed both upregulated and unchanged gene expressions in downstream genes as anticipated from Lysine-Specific Demethylase (LSD1, encoded by *KDM1A*) activity (**Figure 4J**). The results revealed that LSD1, along with several downstream genes including *DOCK9*, *UBR2*, and *ENOX2*, was specifically upregulated. This is the same pattern of modulation neuronal differentiation that has been found for LSD1 (Laurent et al., 2015), which was specifically upregulated in the finch Purkinje cell SVIL+ subtype, exhibiting a notable co-expression pattern. Moreover, gene co-regulation network analysis uncovered a more complex connection network within zebra finch Purkinje cells, suggesting the activation of additional regulatory pathways (**Figure S4H**).

Collectively, cerebellar cell types exhibit significant interspecies variability, suggesting substantial functional evolution within the cerebellum of amniotes, likely in response to diverse lifestyle adaptations. Specifically, it is hypothesized that ALDOC+ Purkinje cells emerged before the common ancestor of amniotes and subsequently diverged across avian and mammalian lineages. In mammals, PLCB4+ Purkinje subtypes developed, whereas in birds, SVIL+ Purkinje subtypes evolved. SVIL+ cells express numerous species-specific marker genes, suggesting roles in dendritic branching and spine morphology. These features are likely crucial for adaptations associated with avian flight.

### Conservation and divergence of non-excitatory neurons in birds

In both individual species and cross-species integration atlas, we found a population of non-excitatory neurons (**Figures 1C and 2A**). These include well-defined definitions derived from medial, lateral, and caudal ganglionic eminences (MGE, LGE, and CGE, respectively) inhibitory neurons (IN), and a cluster of cells with high specific expression of *SNCG*. To investigate evolutionary cell type patterns, we isolated 51,024 non-excitatory cells. Integrated analysis revealed consistent species conservation and co-clustering across species at the level of major cell types (**Figures 5A and 5B**). However, the SNCG+ cell groups, upregulated with *SNCG*, *NEFL*, and *NEFM*, showed the greatest divergence in both the pattern of marker gene expression and the proportion of species represented (**Figures 5C)**. In addition, excitatory characteristics (*SLC17A6* and *SLC17A7*) and inhibitory neuronal characteristics (*GAD1* and *GAD2*) of these cell types were not significant. SNCG+ cell groups were mainly sampled from the hypothalamus of each species and the optic tectum (OT) of pigeons. We analyzed the spatial transcriptomic data from macaques (Lei et al., 2024), zebra finches, and pigeons (Liao et al., 2024) to validate the derived brain regions (**Figure 5D**). According to the results of spatial transcriptomic, the genes expression of SNCG+ cluster (*NEFL* and *NEFM*) in the hypothalamus region was consistent with the characteristics of AVP neurons distributed in the PVN nucleus (Lei *et al*., 2024)(**Figure 5D**). The SNCG+ cell types in bird OT were only distributed in the stratum griseum centrale (SGC) and pars parvocellularis (Ipc) regions (**Figure 5D**). These two nucleus isthmi have been verified in finch to be able to onnected with retinorecipient structures by interneuron (Schmidt and Bischof, 2001), the correlation between their features has worthy of further attention.

**Figure 5.**
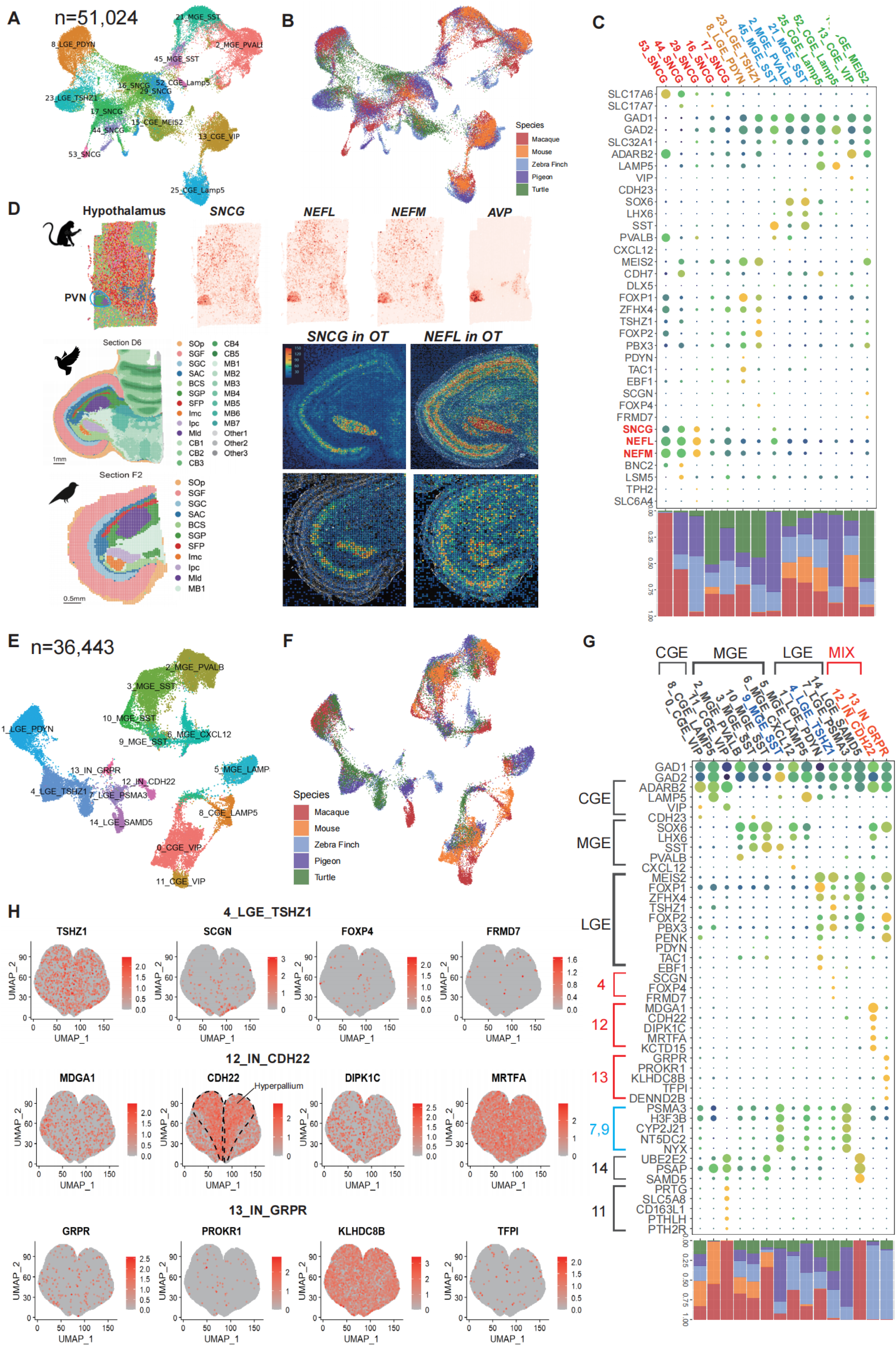
Non-excitatory neurons differentiation across species. **(A-B)** Clustered UMAP plot showing the cell annotation results of the non-excitatory cells containing SNCG from 5 species. The species origin distribution is shown in the (B). **(C)** Dotplots representation of marker genes in each cell cluster in (A). Species origin proportions are identified below with bar plots. The three genes specifically co-expressed by SNCG are highlighted in red. **(D)** Expression of SNCG signature genes in monkey hypothalamus and bird (optic tectal) OT region. **(E-F)** Clustered UMAP plot showing the cell annotation results of the IN cells from 5 species. The species origin distribution is shown in the (F). **(G)** Dotplots representation of marker genes in each cell cluster in (E). Species origin proportions are identified below with bar plots. **(H)** Gene expression on the spatial transcriptome, showing birds-specific enriched marker genes for C4, C12 and C13 in (G).

GABAergic neurons (inhibitory cells, IN), which are more conserved across species such as mice, macaques, and humans than glutamatergic neurons, exhibit less regional brain specificity (Chen et al., 2023; Wang et al., 2022). This conservation is likely due to functional stability. It remains to be determined if this trait persists across a broader range of species. We analyzed 36,443 IN cells, and identified 15 subtypes of IN cells (**Figures 5E and 5F**), originating developmentally from the CGE (c0, c8, and c11), MGE (c2, c3, c5, c6, c9, and c10), LGE (c1, c4, c7, and c14). Cell types of MGE origin were the most abundant among amniotes, encompassing six subtypes and comprising 45% of the cells. Furthermore, MGE-derived cells exhibited the highest conservation of proportion across species. CGE-derived IN cells included two mammal-specific subtypes, 8_CGE_LAMP5 and 11_CGE_VIP, whereas LGE-derived IN cells, such as 14_LGE_SAMD5, were also specific to mammals. All three mammal-specific IN subtypes were characterized by upregulated genes including *UBE2E2*, *PSAP*, and *SAMD5*, associated with cellular lipid metabolism and basal processes. The 4_LGE_TSHZ1 and 7_LGE_PSMA3 subtypes were prevalent among sauropsids and played a significant role in neurodevelopment.

Notably, we identified two cell subtypes with complex origins, 12_IN_CDH22 and 13_IN_GRPR, which were particularly prevalent in sauropsids and abundant in birds (**Figures 5E and 5G**). Subtype 12_IN_CDH22, characterized by the expression of *MDGA1*, *CDH22*, and *KCTD15*, was associated with genes potentially involved in synapse formation and neuronal communication. *CDH22* was predominantly expressed in the hyperpallium of finches (**Figure 5H**), a telencephalic region considered homologous to the neocortex in mammals (Stacho et al., 2020). However, 13_IN_GRPR exhibited a widespread spatial distribution in the finch brain region (**Figure 5H**), expressing *GRPR*, a neuropeptide-related gene, and notably upregulated *FOXP2*, associated with language or speech in humans, and *PENK*, crucial for neural communication (Enard et al., 2002; Haesler et al., 2004; Staes et al., 2017; Sun et al., 2024). Consequently, these abundantly present IN cells in birds may have played a significant role in the functional evolution and adaptive processes of the avian brain.

## Discussion

We explored cell type diversification in the amniote brain, focusing on differences between sauropsids (birds and reptiles) and mammals (primates and rodents) species via single-cell transcriptome analyses. Our study began with a comprehensive single-cell atlas, incorporating approximately 1.3 million cells from the telencephalon and cerebellum of turtles, zebra finches, pigeons, mice, and macaques (**Figure 1A**; **Table S1**). This investigation revealed species-specific variations across a broad range of cell types and subtypes, closely tied to the conservation and diversification of genomes across species. Notably, we observed significant diversification in telencephalic excitatory neurons and various cerebellar cell types between birds and mammals. For instance, most telencephalic excitatory neurons in birds express *SLC17A6*, while mammals exhibit greater regional heterogeneity with *SLC17A7* predominantly in the neocortex and *SLC17A6* in other areas (**Figures 2D and 3E**).

Furthermore, we identified a novel subcell-type of Purkinje cells (SVIL+) in birds (**Figure 4G**), associated with learning and circadian rhythms, featuring numerous positively selected genes in bird-specific Purkinje cell gene expression patterns, underscoring evolutionary adaptations to ecological and behavioral needs (**Figure 4H**). Additionally, we utilized single-cell resolution spatial transcriptomics (Stereo-seq) to analyze the spatial distribution of genes and cell types across amniotes, effectively validating our findings. Collectively, our results establish a robust connection between genetic evolution and environmental adaptation, illustrating how genetic alterations enhance cell type diversification to meet diverse ecological challenges and evolutionary demands.

More specifically, how have the cell types evolved in relation to the species adaptability? In our observations, we identified commonly cell type-specific high expression marker genes exhibiting homologous gene phenomena across various cell types. Notably, *SLC17A6* was highly expressed across the entire pallium in birds, *SLC17A7* in mammals, and both genes were expressed in reptiles (**Figure 2D**). In birds, the transcription factor *NPAS2* in Purkinje cells is co-regulated with *CLOCK*, activating the circadian signaling pathway and facilitating adaptation to environmental changes (**Figures 4H and S4G**). We also observed homologous expression variations of *GABRA4, GABRA6,* and *SOX1, SOX2, SOX9* in granule cells and Bergmann cells (**Figures S4D and S4E**).

Phylogenetic analysis suggests that different expression patterns may arise from two theoretical frameworks: (1) The Gene Balance Hypothesis (Birchler and Veitia, 2010) posits that sequence mutations following genome duplication lead to some copies retaining key life-stabilizing domains while others lose functionality, resulting in paralogous genes expressed distinct patterns across species (e.g. *SLC17A7* in birds). (2) The Duplication-Degeneration-Complementation Model (Force et al., 1999) suggests that homologous genes subfunctionalize, share ancestral functions, and perform these functions through dose compensation, often seen with multiple homologous genes expressing concurrently (e.g. *SLC17A7* and *SLC17A6* in turtles).

Similar findings in comparative studies of the hypothalamus indicate that various species regulate cell functions through subfunctionalization and dose-compensation effects of homologous genes (Shafer *et al*., 2022). Our study extends these conclusions across amniotes and down to specific gene domains, such as mutation in *SLC17A7* in zebra finch, leading to loss of a key neurotransmitter transmission function and compensatory expression of paralogous *SLC17A6* (**Figure 3G**). This adaptive expression is observed in various cerebellar cell types as well (**Figures S4D, and S4E**). Furthermore, sequence differentiation results in the retention of differentially expressed genes that positively impact survival, highlighting that positively selected genes exhibit the most detailed species differences. Conversely, transcription factors (TFs) are more conserved due to their crucial role in influencing downstream gene expression (**Figure S2D**).

From a comprehensive perspective, gene duplication, mutation, and loss events in genome evolution significantly influences the cross-species evolution of cell types. Transcription factor (TF) genes demonstrate greater stability due to the universality of their functions (Weidemuller et al., 2021). We propose that ancestral key genes underwent duplication events, resulting in multiple homologous genes that may play a more important role in the cell type evolution. These genes were retained in the genomes of different species during species divergence. They underwent gains and losses of gene motifs, and even complete gene deletions, which significantly influenced the differences in the gene regulatory network, ultimately affecting the differentiation and distribution of cell types.

In the end, identifying and recognizing homologous cell types is a pivotal challenge in conducting cross-species single-cell studies to explore the evolution of cell types, structures, and functions. In this study, we employed a gene family homology approach, instead of solely relying on one-to-one orthologous gene information as typically done. This method enabled us to connect homologous genes across distantly related species on a larger scale, expanding the analysis to thousands of gene families. While this approach mitigates issues related to paralogous relationships, it might overlook functional differences in paralogous genes, a limitation that future research needs to address. Furthermore, we adopted an enhanced method based on Graph Convolutional Networks (GCN), termed entropy-scGCN, which effectively integrates single-cell data from multiple species using canonical marker information, providing a solid basis for identifying homologous cell types. With the increasing application of large language models in life sciences, particularly for cross-species comparisons such as SATURN (Rosen et al., 2024) and UCE (Rosen et al., 2023), there is significant promise for future advancements. As more high-quality single-cell data from various species become available, employing large models in cross-species studies is expected to substantially improve our understanding of homologous cell types and their evolutionary trajectories.

## Author contributions

S.L., R.N., X.X., Z.W., C.X., D.C. conceived and designed the study. D.C., S.L., R.N., Zhenkun.Z, wrote the manuscript and F.H., Y.W., K.L., H.W., J.H., Ying L., Y.B., C.X., J.W., H.Y., Yidi S., G.C., M.P., Yangang S., Ze.Z., L.L., Youning L., contributed to the discussion and revision of the manuscript. H.W., Z.W., M.H., W.X. contributed to sample collection. T. Zeng, Y.D. contributed to the sequencing. Y.H, M.C., J.P. designed and performed snRNA-seq and stereo-seq, D.C., Zhenkun.Z., Y.Y., Y.Z., F.H., K.L., X.J., Y.X., Z.N., Zhiyong.Z., discussed and conducted data analysis. X.W. and T.Y. collected the analyzed data and established the online database. All authors read and approved the final manuscript.

## Acknowledgments

This paper is from the Mesoscopic Brain Mapping Consortium. The project was supported by National Key R&D Program of China (2022YFC3400405, 2021YFA0805100, and 2020YFE0205900), National Science and Technology Innovation 2030 Major Program (STI2030-2021ZD0200100, STI2030-2022ZD0205000 and STI2030-2022ZD0211700), Natural Science Foundation of Guangdong Province, China (2021B1515120075) and Guangdong Genomics Data Center (2021B1212100001), Zhejiang Province ’Vanguard’ R&D Program (2023C03SA103409), and Hangzhou Leading Innovation Team Project. We sincerely thank the China National GeneBank for providing technical support.

## Declaration of generative AI and AI-assisted technologies in the writing process

During the preparation of this work the authors used ChatGPT to improve the language. After using this tool, the authors reviewed and edited the content as needed and took full responsibility for the content of the published article.

## RESOURCE AVAILABILITY

**Lead contact**

Further information and requests for the resources and reagents may be directed to and will be fulfilled by the lead contact, Shiping Liu.

## Materials availability

All materials used for stereo-seq and snRNA-seq are commercially available.

## Data and code availability

The data that support the findings of this study have been deposited into CNGB Sequence Archive (CNSA) (Guo et al., 2020) of China National GeneBank DataBase (CNGBdb) (Chen et al., 2020) with accession number CNP0003026 (https://db.cngb.org/search/project/CNP0003026/). Additionally, processed stereo-seq and snRNA-seq data used in this study can be accessed and downloaded via STOmicsDB (Xu et al., 2024): https://db.cngb.org/cdcp/dataset/SCDS0000639/. The updated and improved entropy-scGCN based on scGCN algorithm is available on GitHub (https://github.com/Dee-chen/entropy-scGCN/)All data were publicly available as of the date of publication. Any additional information required to re-analyze the data reported in this paper is available from the lead contact upon request.

## EXPERIMENTAL MODEL AND SUBJECT DETAILS

### Animals

Experimental procedures were performed in accordance with the guidelines of the Animal Care and Use Committees at the Shenzhen Institute of Advanced Technology (SIAT), Chinese Academy of Sciences (CAS), China (permit number SIAT-IACUC-210326-NS-WH-A1881). Experiments were performed using 8 male Chinese softshell turtles (Pelodiscus sinensis) weighing 450-600 g, 8 female and male adult (>12 months old) zebra finches (Taeniopygia guttata), and 2 pigeons (Columba livia) at 12 months of age. Turtles were obtained from external breeders and zebra finches from the colony at Zhengzhou university, China.

## METHOD DETAILS

### Library preparation and sequencing

One-hundred ng of cDNA (20 μl) from each sample were tagmented with Tn5 transposases (Vazyme) at 55C for 10 mins, followed by quenching the reaction using 5 μL of 0.02% SDS. Subsequently, a PCR reaction mix (75 μL, Library HIFI Master Mix, Library PCR primer mix) was added to each fragmented cDNA sample. The samples were then subjected to thermal cycling for amplification using the following protocol: an initial cycle at 95C for 5 min, followed by 13 cycles of tri-temperature reactions (98C for 20 s, 58C for 20 s and 72C for 30 s), and a final cycle at 72C for another five minutes. After amplification, the PCR products were purified using VAHTSTM DNA Clean Beads (VAZYME, N411-03) in two steps with concentrations of beads set as follows: first step with a ratio of bead volume to sample volume as low as approximately six-tenths and second step with a ratio around two-tenths; subsequently these purified products were utilized in DNB (DNA Nano Ball) generation process. Finally, the generated DNBs underwent sequencing on the DNBSEQ^TM^ T10 platform (MGI, Shenzhen, China), employing read1 length of fifty base pairs and read2 length of one hundred base pairs.

### Brain tissue collection for snRNA-seq

The snRNA-seq samples were collected from frozen sections adjacent to those for Stereo-seq. These sections were cut at 50-μm thickness, and 3 to 5 sections for each coronal coordinate were collected for snRNA-seq analysis. Sections were transferred into plastic wells on dry ice and stored in a -80 C refrigerator. Each section was further segmented into distinct areas on dry ice using tissue punchers (5 - 8 mm in diameter). Tissues at the same brain regions were combined in a prechilled pipe as one sample. In the cryostat, the cortical areas were segmented at 1 - 2 mm depth using tissue punchers (2.5 - 4 mm in diameter). After dissection, the samples were immediately frozen in liquid nitrogen and then kept in dry ice or -80 C refrigerator. Throughout the sampling manipulation, the tissues were carefully transferred to pre-cold tube without thawing.

### Single-nucleus suspension preparation

Single nucleus suspension was prepared as previously described (Bakken et al., 2018). Briefly, frozen brain tissue pieces were placed in Dounce homogenizer with 2 ml pre-chilled homogenization buffer and kept the Dounce homogenizer on ice during grinding. Tissue was homogenized with 10-15 strokes of the pestle A and followed by 10-15 strokes of the pestle B, then added 2 ml homogenization buffer to the Dounce homogenizer and filtered the homogenate through 30μm MACS SmartStrainers (Miltenyi Biotech, #130-110-915) into 15 ml conical tube and centrifuged at 500 g for 5 mins at 4C to pellet nuclei, then the pellet was resuspended in 1.5 ml of blocking buffer and centrifuged at 500 g for 5 mins at 4C to pellet nuclei. Nuclei were resuspended with cell resuspension buffer for subsequent snRNA-seq library preparation.

### snRNA-seq data processing

Initially, bead barcodes and unique molecular identifier (UMI) sequences were extracted using the parse function in PISA (https://github.com/shiquan/PISA). For cDNA libraries, read1 encompassed bead barcodes from positions 1-10bp and 11-20bp, while the UMI sequence was located at position 21-30bp. The entire read2 (100bp) was utilized for downstream alignment analysis. In the case of Droplet Index libraries, read1 contained bead barcodes at positions 1-10bp and 11-20bp, with the UMI sequence present at position 1-10bp of read2. Additionally, droplet index barcodes were found at positions 11-20bp and 21-30bp of read2. Reads with incorrect barcodes based on the barcode list were excluded from further analysis. Subsequently, STAR (Dobin et al., 2013) was employed to align snRNA-seq data against the genome reference. To estimate the actual number of beads, we used the “barcodeRanks” function of DropletUtils tool (https://github.com/MarioniLab/DropletUtils/) to find the threshold value of sharp transition in total UMI counts distribution. Beads with UMI counts less than the threshold were removed. We merged the beads considered to be one cell, and counted the gene expression of cells by PISA.

### Gene family clustering and evolutionary event identification

In order to determine the homology relationship of gene sequences of amniotic animals and locate evolutionary events such as gene duplication, acquisition and loss, and positive selection, we downloaded CDS and protein sequences of mice, humans, cynomolgus monkeys, and zebra finches from NCBI data. We used third-generation sequencing to re-annotate existing gene annotations for pigeons and Chinese soft-shell turtles. After that, we performed gene family clustering by Tree2gd (Chen et al., 2022b) on zebrafish with sequence of each species for the outgroup, and a total of 38,948 gene families were obtained. Tree2gd was used to locate the evolutionary event and calculate the Ka/Ks of homologous genes.

### Cell downsampling and cross-species integration

Basic processing and visualization of the scRNA-seq data were performed with the SeuratCv.4.3.0C in RCv.4.2.2C.We discarded cells with the number of genes (nFeatureRNA) less than 2,000, the number of genes (nCount) more than 2,500 and the percentage of mitochondrial genes (percent.mt) larger than 5%.After the first quality control, 195,162 cells were remained for the following analyses.We integrated the data from mice, macaque, pigeons, zebra finches, and turtles using the “IntegrateData” function of the Seurat R package (v 4.3.0), with marker genes as anchors.Then principal component analysis (PCA) was carried out and the top 30 components were used for downstream analysis. The integrated datasets were re-clustered with k.param = 20 and resolution = 0.5 using “FindNeighbors” and “FindClusters” functions.We then categorized the 63 cell clusters into 19 categories based on classical marker genes, including glutamatergic neurons, GABAergic neurons and non-neuronal cells. Then the 19 categories of cells were further iteratively classified into more cell clusters with higher resolution.

### Update turtle and pigeon genome annotations by IsoSeq

The PacBio data were processed using IsoSeq v3.4.0 (https://github.com/PacificBiosciences/IsoSeq). Only sequence reads containing both 5’ and 3’ adaptors were retained to cover the entire transcript. Subsequently, the corrected SMRT reads were aligned to the reference genome PelSin_1.0 (Wang et al., 2013) and Cliv_2.1 (Shapiro et al., 2013) using GMAP (Wu et al., 2016) to locate the position of the predicted genes on the chromosomes.A total of 60,167 and 48,142 transcripts were successfully aligned to the genome. We used Gffcompare (https://ccb.jhu.edu/software/stringtie/gffcompare.shtml) (Pertea and Pertea, 2020) to compare the assembled transcript models with the transcript models of the reference genome and concordant transcripts (class code “=”) were merged. For cross-species comparisons, the combined annotated protein sequences were aligned using BLAST+ (Camacho et al., 2009) and proteins from other species in the article. The genes on the alignment were renamed to their corresponding gene names, and the others retained the gene ID given by the software. The snRNA-seq ratio was significantly improved in the data in this paper (table below). The gtf file is available at https://github.com/Dee-chen/Refined-gtf-for-turtle-and-pigeon.

| Species | Turtle |  | Pigeon |  |
| --- | --- | --- | --- | --- |
| Gtf version | PelSin_1.0 | Refined gtf | Cliv_2.1 | Refined gtf |
| Reads mapped to genome | 88.50% | 88.50% | 91.00% | 91.00% |
| Reads mapped to exonic regions | 18.30% | 27.90% | 26.60% | 35.80% |
| Reads mapped to intronic regions | 28.60% | 31.90% | 21.80% | 40.40% |
| Reads mapped across exon and intron regions | 0.90% | 1.00% | 1.70% | 1.70% |
| Reads mapped antisense to gene | 14.20% | 15.40% | 6.70% | 7.80% |
| Reads mapped to intergenic regions | 37.80% | 23.70% | 42.90% | 13.90% |
| Unidentified type | 0.10% | 0.20% | 0.30% | 0.30% |

### Conservative entropy calculation across species cell types

We have innovated and improved on scGCN (Song *et al*., 2021) by developing a multi-omics cross-species cell-type annotation and conservation assessment software sc-Entropy based on information entropy theory. Firstly, the FASTA format protein sequence files of the desired species were downloaded, and subsequently the Tree2GD (Chen *et al*., 2022b) tool was used to identify the gain, loss, and duplication events of all genes based on the relationship between the sequences to construct the gene family evolutionary lineage tree. Therefore, entropy-scGCN is not only suitable for cell type mapping of single genes, but also for cell type mapping at the gene family level, solving the limitation of using only one-to-one homologous genes. In order to measure how conservative the prediction results of different subgroups are, we introduce the calculation of information entropy in entropy-scGCN. entropy-scGCN was used to obtain the percentage P_ij_ of each subgroup in the test data set predicted to be the cell type in the reference data set. The basic information entropy could be obtained by using the calculation formula of information entropy (see Equation 1).

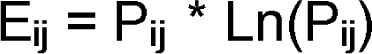

The original entropy value only contains information about whether the probability distribution of mapped cell types obtained by entropy-scGCN is reasonable, without considering the influence of the similarity relationship between cell types in the reference data set. In order to reduce the influence caused by the differences between cell types in the reference data set, MetaNeighbor (Fischer et al., 2021) was used to calculate the similarity of cell types in the reference data set, and the similarity score between cell types was obtained (Dis_j_). The matrix multiplication of the similarity score with the basic information entropy was performed, and the corrected information entropy was obtained by normalization. The smaller the entropy value, the more homogeneous the cell type obtained by entropy-scGCN annotation, that is, a higher proportion of the cell type was annotated as one cell type rather than predicted as multiple cell types. The formula for calculating entropy value after adding the distance factor between cell types in the reference data set is as follows (see Equation 2):

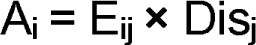

### Cell type differentially expressed gene analysis and species comparison

In this study, the data was split by species, and differential gene expression analysis was performed using the “FindAllMarkers” function. To identify genes that were significantly differentially expressed, we applied a filter with the criterion avg_log2FC > 1, resulting in a range of 43,864 to 56,490 differentially expressed genes.To investigate the commonalities among the different species, we assessed the intersection of these differentially expressed genes. The resulting gene intersections were visualized using UpSetR v1.4.0, and an UpSet plot was generated to illustrate the overlapping gene sets.

Furthermore, we extended our analysis to gene families by following a similar approach. We obtained gene families for each species, encompassing a range of 34,151 to 51,547 gene families. To identify the intersections of gene families among the different species, we employed UpSetR v1.4.0.

## Supplementary Figure Legend

**Figure S1.**
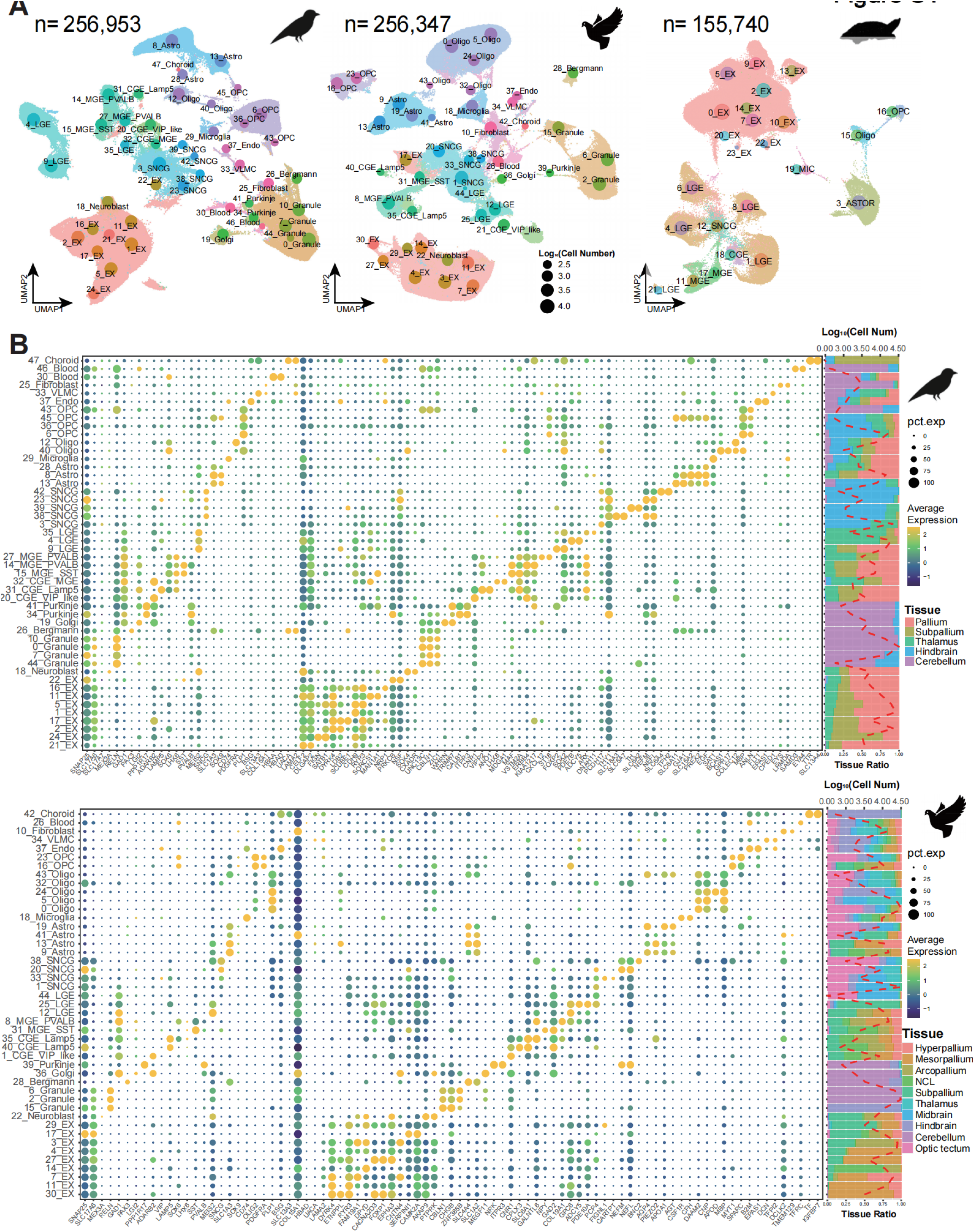
snRNA-seq atlas of sub cell-types in zebra finch, pigeon, and turtle. **(A)** UMAP plots of snRNA-seq atlas of zebra finch, pigeon, and turtle, colored by sub cell-types, and the size of the point indicated the number of cells. **(B)** Dot plot showing the expression of key marker genes for cell subclasses in zebra finch (upper panel) and pigeon (lower panel). Marker genes are categorized into major cell class marker genes (front) and sub cell-type DEGs. A histogram on the right shows the proportion of brain regions sampled, while the red line represents the logarithm of the number of cells.

**Figure S2.**
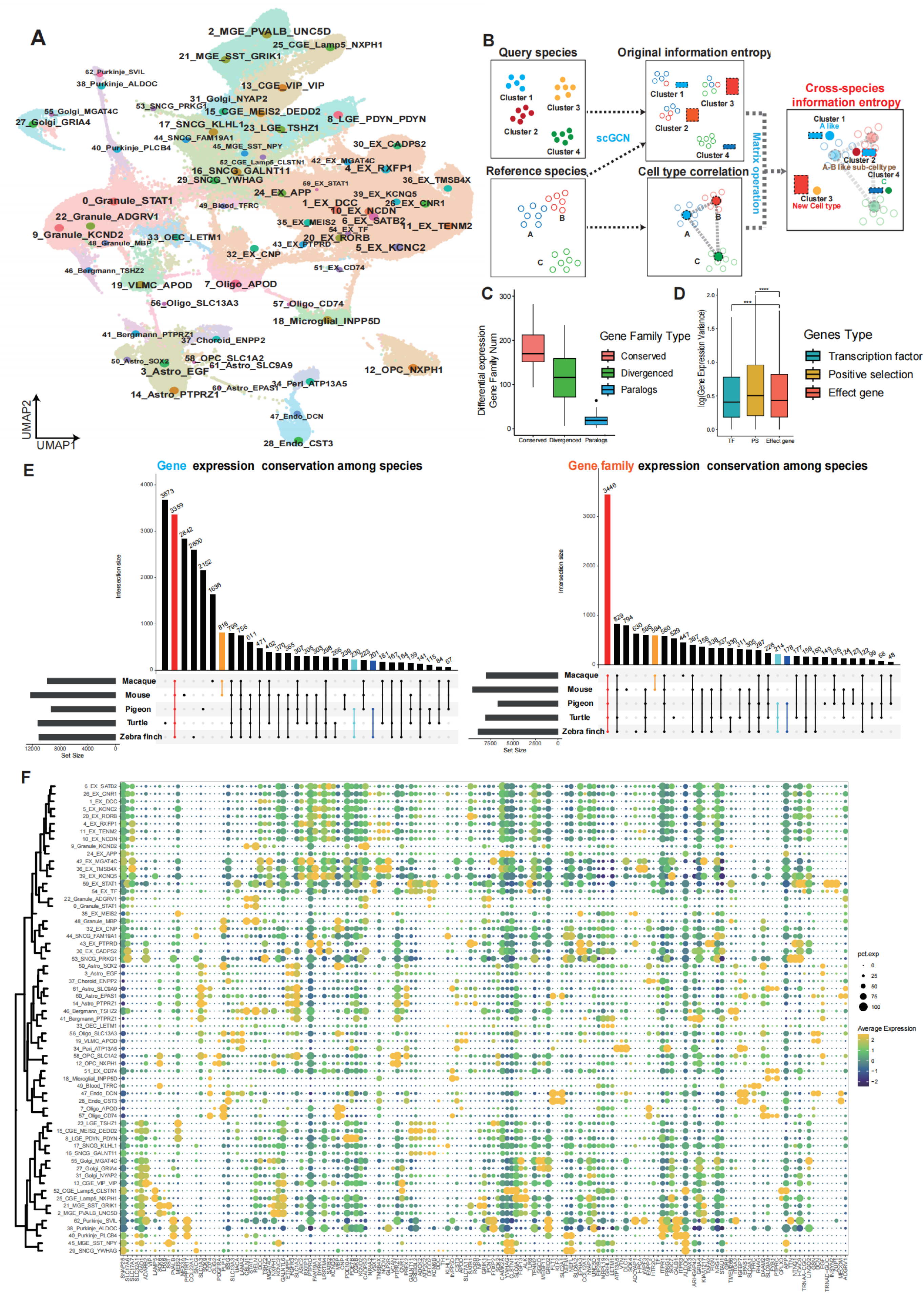
Comparison of gene expression across species cell types. **(A)** Clustering UMAP of 63 sub cell-types of 5 species integration corresponding to Figure 2A and D. **(B)** Schematic diagram of the calculation of “cross-species conservation information entropy”, an evaluation index of cell type conservation degree in entropy-scGCN. **(C)** Box plot of the number of DEGs in each cell type, which classified by the sequence conserved genes (red), sequence differentiated genes (green) and paralogs genes caused by gene duplication (blue). **(D)** Box plot shows the log differential expression multiples of various genes in the same cell type between species, and the genes are divided into transcription factors (TF, blue), positive selection genes (PS, red), and other effect genes (gray) according to their characteristics. **(E)** The upset plot displays the distribution of differentially expressed genes (DEGs) and differentially expressed gene families (DEGFs) across all cell types of five species in (A). Evolutionarily conserved types are highlighted with colors (red: amniotes; yellow: mammals; teal: squamates; blue: birds) **(F)** Dot plot of the expression of top3 differentially expressed genes in 63 cell types.

**Figure S3.**
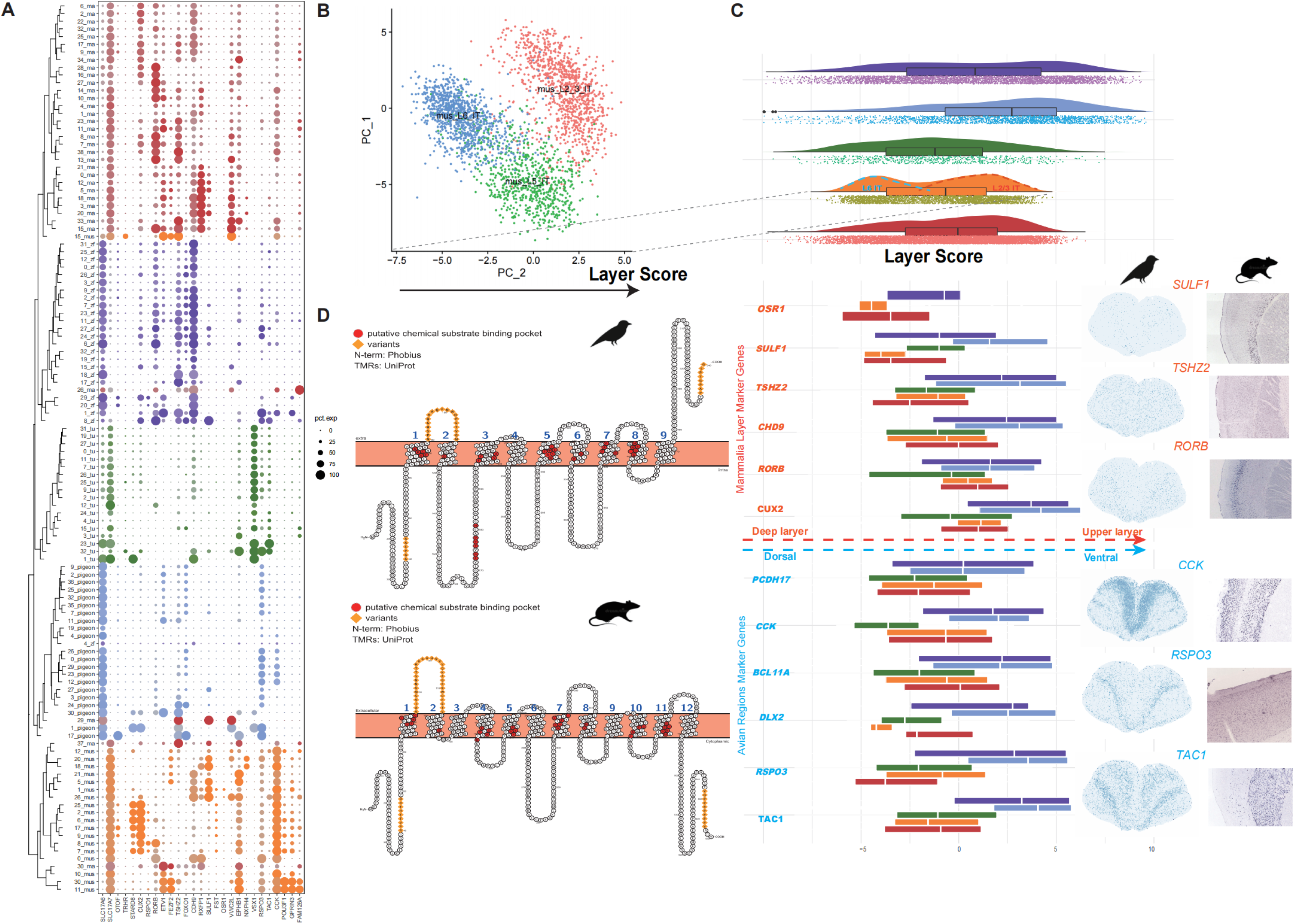
Comparison of differences in excitatory neurons across species. **(A)** Dot plot of species-split cortical excitatory neuron marker genes. **(B)** PCA cluster map of excitatory neurons in mouse cortex. PC2 is the principal component that can maximally distinguish the Up layer from the Deep layer. The gene weight of PC2 was set as the “Laryer Score”, which was applied in the excitability clustering with other species. **(C)** 5 species layer score distribution and key gene expression. The above shows the violin diagram of the overall distribution of layer score of different species of excitatory neurons. The bottom part shows the expression distribution of key genes in layer score. The red and blue are the key genes in mammals and birds, respectively. The expression of key genes in space is shown on the right. **(D)** Schematic representation of transmembrane structure and sequence of SLC17A7 gene in mouse and zebra finch. Red amino acids are key binding pockets, and yellow is the fragment where the protein sequence is mutated between the two species.

**Figure S4.**
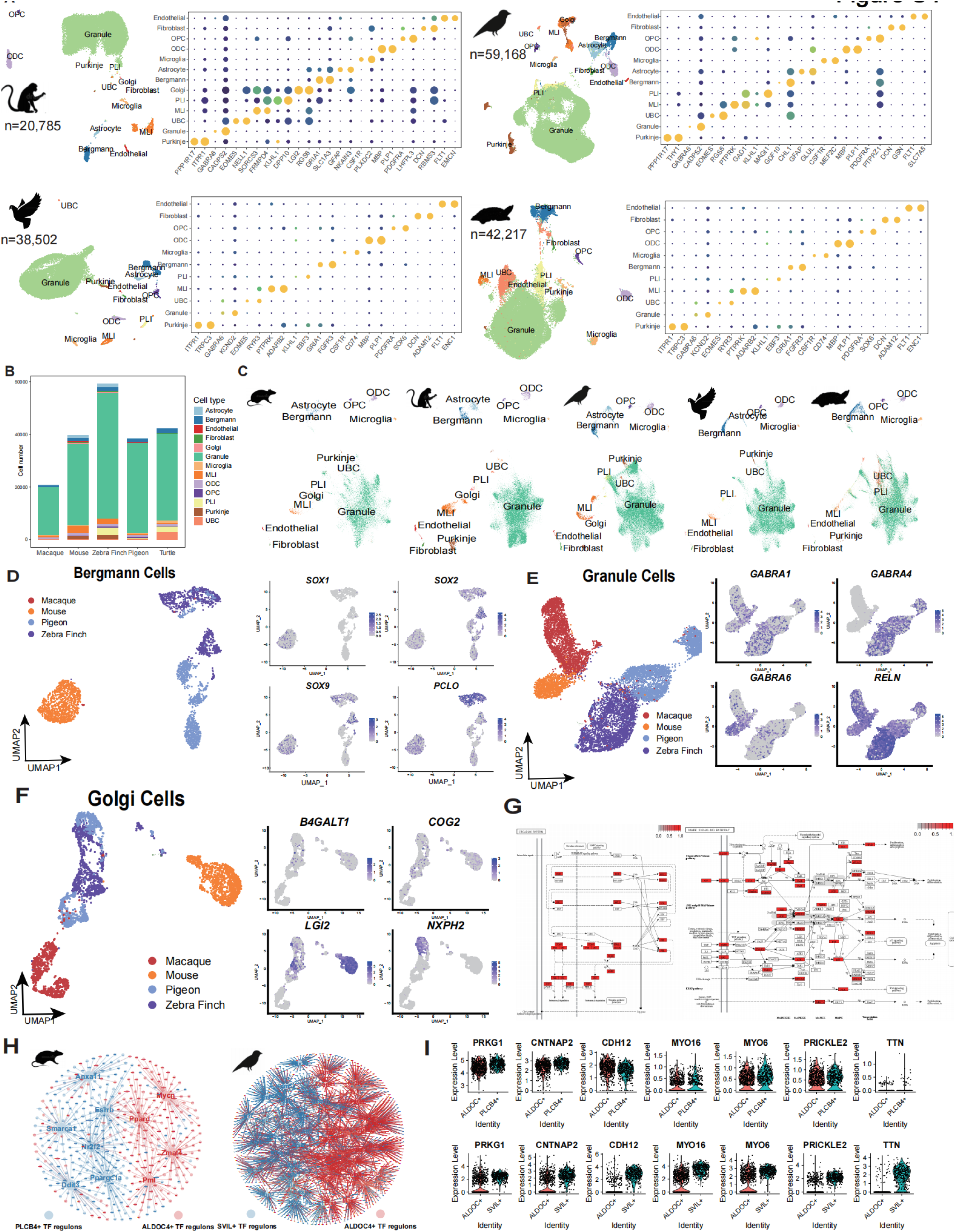
Comparison of differences in cerebellum cell types across species. **(A)** UMAP of clustering of cerebellar cell types and marker gene expression plots of the four species. **(B)** Bar plot of proportional distribution of cerebellar cell types in the five species. **(C)** UMAP plot of different species origin split in cerebellar cell type integration (Figure4A). **(D-F)** Bergmann cells, granule cells, and Golgi cells were clustered separately for UMAP maps and key paralogous marker gene expression. **(G)** KEGG pathway proteins for circadian rhythms and MAPK. Genes highly expressed in zebra finch SVIL+ are shown in red. **(H)** Differentially expressed TF regulon between mouse and zebra Purkinje cell subtypes obtained by SCENIC (Van de Sande et al., 2020). **(I)** Violin plot of differential expression of PIP2 hydrolysis pathway genes in mouse and zebra finch Purkinje cells

## Supplementary Table

**Table S1. Single cell atlas cell classification and number of downsampled cells, related to** Figure 1 **and** Figure 2

(1-5) Data set statistics for pigeons, zebra finches, monkeys, mice, and turtles

**Table S2. The result table of gene family correspondence between species is obtained by Tree2GD, related to Figure 2 and Figure S2**

**Table S3. Differentially expressed genes (DEGs) and corresponding differentially expressed gene families (DEGFs) of each 63 cell types split by species, related to** Figure 2 **and Figure S2**

(1-5) DEGs and DEGFs statistics for pigeons, zebra finches, monkeys, mice, and turtles

**Table S4. Using mice as reference, the entropy values of 63 cell types of the remaining 4 species obtained by Entropy-scGCN, related to** Figure 2

**Table S5. List of transcription factors (TFs) used in the study, related to Figure S2 and** Figure 4

**Table S6. List of Positive selection genes (PSGs) used in the study, related to Figure S2 and** Figure 4

**Table S7. KEGG enrichment of mouse and zebra Purkinje cell category DEGs, related to** Figure 4

